# Applications of community detection algorithms to large biological datasets

**DOI:** 10.1101/547570

**Authors:** Itamar Kanter, Gur Yaari, Tomer Kalisky

## Abstract

Recent advances in data acquiring technologies in biology have led to major challenges in mining relevant information from large datasets. For example, single-cell RNA sequencing technologies are producing expression and sequence information from tens of thousands of cells in every single experiment. A common task in analyzing biological data is to cluster samples or features (e.g. genes) into groups sharing common characteristics. This is an NP-hard problem for which numerous heuristic algorithms have been developed. However, in many cases, the clusters created by these algorithms do not reflect biological reality. To overcome this, a Networks Based Clustering (NBC) approach was recently proposed, by which the samples or genes in the dataset are first mapped to a network and then community detection (CD) algorithms are used to identify clusters of nodes.

Here, we created an open and flexible python-based toolkit for NBC that enables easy and accessible network construction and community detection. We then tested the applicability of NBC for identifying clusters of cells or genes from previously published large-scale single-cell and bulk RNA-seq datasets.

We show that NBC can be used to accurately and efficiently analyze large-scale datasets of RNA sequencing experiments.

## Introduction

Advances in high-throughput genomic technologies have revolutionized the way biological data is being acquired. Technologies like DNA sequencing (DNA-seq), RNA sequencing (RNA-seq), chromatin immunoprecipitation sequencing (ChIP-seq), and mass cytometry are becoming standard components of modern biological research. The majority of these datasets are publicly available for further large-scale studies. Notable examples include the Genotype-Tissue Expression (GTEx) project^1^, the cancer genome atlas (TCGA)^2^, and the 1000 genomes project^3^. Examples of utilizing these datasets include studying allele-specific expression across tissues^4, 5^, characterizing functional variation in the human genome^6^, finding patterns of transcriptome variations across individuals and tissues^7^, and characterizing the global mutational landscape of cancer^8^. Moreover, some of these genomic technologies have recently been adapted to work at the single-cell level^9^. While pioneering single-cell RNA sequencing (scRNA-seq) studies were able to process relatively small numbers of cells (42 cells in^10^ and 18 cells in^11^), recent single-cell RNA-seq studies taking advantage of automation and nanotechnology were able to produce expression and sequence data from many thousands of individual cells (*∼*1,500 cells in^12^ and *∼*40,000 cells in^13^). Hence, biology is facing significant challenges in handling and analyzing large complex datasets^14, 15^.

### Clustering analysis

One of the common methods used for making sense of large biological datasets is cluster analysis: the task of grouping similar samples or features^16^. For example, clustering analysis has been used to identify subtypes of breast tumors^17, 18^ with implications to treatment and prognosis. More recently, clustering analysis was used to identify and characterize cell types in various tissues and tumors in the colon^19^, brain^20^, blood^12^, and lung^21^, with the overall aim of finding key stem and progenitor cell populations involved in tissue development, repair, and tumorigenesis. Another application is to find sets of coordinately regulated genes in order to find gene modules^11, 22, 23^. Such clusters of genes (or other features such as single-nucleotide polymorphism (SNPs)^24^) can be further analyzed by gene set enrichment approaches to identify gene annotations^25^ (e.g. GO^26^, KEGG^27^, and OMIM^28^) that are over-represented in a given cluster, and thus shed light on their biological functionalities^29^. Two of the most common clustering methods used in biology are K-means clustering, which groups data-points into *K* prototypes (where *K* is a predetermined number of clusters), and hierarchical clustering, which builds a hierarchy of clusters from all data points^30^. Other methods for clustering include self-organizing map (SOM)^31^, spectral clustering^32^, and density based methods^33^ (for a comprehensive review on clustering see^30, 34, 35^).

### Networks and community detection

Another way to model biological systems is through network science. Networks (also known as graphs) are structures composed of nodes that are connected by edges, which may be weighted and/or directional. Networks have been used for modeling interactions between components of complex systems such as users in social media platforms (Facebook^36^ and Twitter^37^) or proteins^38^ and genes^39^ in a cell. Often, networks contain communities of nodes that are tightly interconnected with each other, which is an indication of common features. For example, communities in social networks^40^ are composed, in most cases, of people with common interests or goals that are interacting with each other. Likewise, proteins with related functionalities interact with each other, and as a result form close-knit communities in protein-protein interactions (PPI) networks^41^. Similarly, web pages with similar content in the World Wide Web^42^ usually have links to each other.

The problem of community detection (CD), which can be viewed as a network’s equivalent for clustering, is not rigorously defined. As a consequence, there are numerous CD algorithms that solve this problem reasonably well using different strategies^43^. An intuitive way to define communities in a network is to divide the nodes into groups that have many in-group edges and few out-group edges. This can be achieved by maximizing the network modularity - a measure that quantifies edge density within communities compared to edge sparseness between communities^44, 45^ (see Methods for formal definition).

Numerous community detection algorithms were developed during the last two decades^43, 46^. Newman defined a measure for the modularity of a weighted network^45^. Clauset et al. developed a fast greedy algorithm to partition the nodes of the network in a way that maximizes the modularity by hierarchical agglomeration of the nodes^47^. Reichardt et al. proposed an approach based on statistical mechanics. Their algorithm models the network as a spin glass and aims to find the partition of nodes with maximal modularity by finding the minimal energy state of the system^48^. Rosvall et al. and Yucel et al. proposed methods to find the community structure by approximating the information flow in the network as a random walk^49, 50^. Jian et al. developed SPICi - a fast clustering algorithm for large biological networks based on expanding clusters from local seeds^51^.

A popular community detection algorithm called the *Louvain* algorithm was proposed by Blondel et al.^52^ (see Methods for details). The *Louvain* algorithm starts by defining each node as a separate community and then performs modularity optimization by an iterative heuristic two-step process. In the first step, the algorithm goes over all nodes of the network and checks, for each individual node, if the network modularity can be increased by removing it from its present community and joining it to one of its neighboring communities. The process is repeated until no further increase in modularity can be achieved. This approach is called the “local moving heuristic”. In the second step, a new meta-network, whose nodes are the communities identified by the first step, is constructed. The two steps are repeated until maximum modularity is attained. It was shown that this algorithm can be improved further by modifying the “local moving heuristic”. SLM, for example, attempts to split each community into sub-communities prior to construction of the meta-network. In this way, communities can be split up and sets of nodes can be moved from one community to another for improving the overall modularity score^53^.

Recently, several networks-based clustering algorithms were developed specifically for single-cell gene expression datasets (see Table 1). Pe’er and colleagues developed PhenoGraph^54^. This method first builds a k-nearest neighbors (KNN) network of cells, where each cell is connected to its *K* nearest cells in Euclidean space. In order to better resolve rare or non-convex cell populations, the algorithm then constructs a second, shared nearest neighbors (SNN) network, in which the similarity between every two nodes is determined by the number of neighboring nodes that are connected to both of them. Finally, the *Louvain* community detection algorithm is used to find groups of cells with similar gene expression profiles. Applying their method to mass cytometry data from 30,000 human bone marrow cells, they were able to cluster single-cell expression profiles into different immune cell types. Su and Xu developed another algorithm called SNN-cliq^55^. This algorithm also constructs a KNN network, and then constructs a SNN network in which the weight between every two nodes is determined not only by the number of shared nearest neighbors, but also their distances to the two nodes. Communities then are detected using a quasi-clique-based clustering algorithm. When applying their method to several single-cell transcriptomics datasets, they found it to be more robust and precise than traditional clustering approaches.

**Table 1.** Examples of previously published methods for NBC.

| Name | Reference | Description |
| --- | --- | --- |
| PhenoGraph | Manuscript:<sup>54</sup><br>Software:<sup>96</sup> | A KNN network is constructed using Euclidean distance. Then, a SNN network is constructed by using the number of shared neighbors between every two nodes as a new similarity measure between them. Communities are found using the *Louvain* algorithm. |
| SNN-Cliq | Manuscript:<sup>55</sup><br>Software:<sup>97</sup> | A KNN network is constructed using Euclidean distance (or similar). Then, a SNN network is constructed using the number of shared neighbors between every two nodes, as well as their distances to the two nodes, as a new similarity measure. Communities are found using a heuristic approach to find "quasi-cliques" and merge them. |
| Seurat | Manuscript:<sup>56</sup><br>Software:<sup>98</sup> | The algorithm constructs a KNN network and then a SNN network. Communities are found using the *Louvain*<sup>52</sup> or the SLM<sup>53</sup> algorithms. |

In this manuscript, we introduce an accessible and flexible python-based toolkit for Networks Based Clustering (NBC) for large-scale RNA-seq datasets in the form of an *IPython* notebook with a self-contained example (Supplementary File S1 online). This toolkit allows the user to follow and modify various aspects of the algorithms detailed above^52–56^, that is, to map a given dataset into a KNN network, to visualize the network, and to perform community detection using a variety of similarity measures and community detection algorithms of his choice. This flexibility is important since, from our experience, different parameters and algorithms might work best for different datasets according to their specific characteristics. Using this toolkit, we tested the performance of NBC on previously published large-scale single-cell and bulk RNA-seq datasets.

## Results

### A workflow for Networks Based Clustering (NBC)

A typical Networks Based Clustering workflow can be divided into four steps (Fig 1). The given dataset, in the form of a matrix of N samples (e.g. cells) by P features (e.g. genes), is first preprocessed by normalizing the samples to each other^57, 58^ and filtering less informative samples or features^58, 59^. This step is especially important in single-cell data since it is typically noisier than bulk samples. Likewise, in meta-analysis, it is important to normalize samples obtained from different sources in order to to mitigate bias due to batch effects^60, 61^. In our *IPython* notebook we took a dataset that was collected and pre-filtered by Patel et al.^62^ and normalized each sample such that it will have zero mean and unit length (L2 normalization).

**Figure 1.**
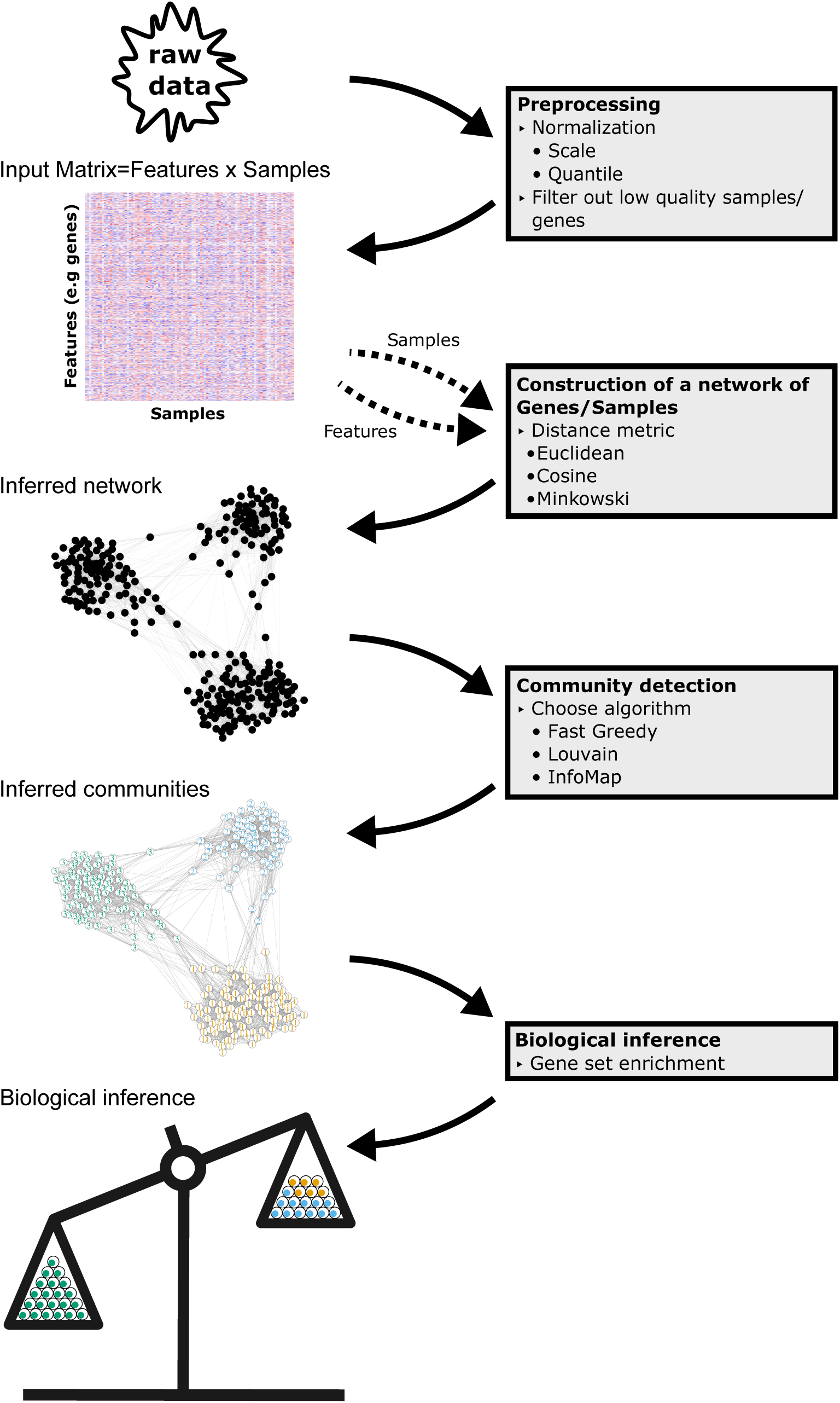
A typical workflow for Networks Based Clustering (NBC). The raw data is first preprocessed to form a gene expression matrix. From this matrix, a weighted KNN network is constructed, in which each node represents a sample (e.g a single cell) or a feature (e.g. a gene). Then, a community detection algorithm is applied to partition the nodes into closely-knit communities, which can be characterized using enrichment strategies.

Next, a similarity measure is defined between the samples (or alternatively, features) and a KNN (K-nearest neighbors) network is constructed as follows: First, each sample (or feature) is represented as a node. Then each node is linked to its *K* nearest neighboring nodes. Constructing a KNN network using the naïve algorithm has a complexity of *O*(*N*^2^) which is slow for large N. We therefore use the more efficient ball tree algorithm^63^, whose complexity scales as *O*(*N* log(*N*)) if supplied with a true distance metric that satisfies the triangle inequality^64^. There are various such distance-like measures (each based on a similarity measures) that can be used^65^ to construct the network, for example, the Euclidean distance, the Minkowski distance, and the cosine distance. In our solved example we used the cosine similarity (see Methods). Note that the popular correlation distance does not satisfy the triangle inequality^64^ and hence is not a true metric. Following network construction, a community detection (CD) algorithm is performed, resulting in an assignment of each node (sample or feature) to a distinct community.

Once communities have been identified, each community can be characterized to infer its biological meaning. For example, communities of cells may represent cell sub-populations in complex tissues or tumors and can be identified using previously known markers^66, 67^. Similarly, the biological functionality of communities of genes (or of gene sets that are over-expressed in specific cell communities) can be inferred using enrichment analysis at the gene and gene-set levels^29, 68–72^.

### NBC accurately resolves seven cell types from a glioblastoma single-cell RNA-seq dataset

To test the performance of NBC, we analyzed single-cell RNA-seq datasets originally published by Patel *et al.*^62^, for which the biological interpretation was known a-priori. These datasets were previously obtained from five glioblastoma patients and two gliomasphere cell lines and were found to contain 7 biologically distinct cell types. A 2D representation of the data by PCA and tSNE can be found in Supplementary File S1 online. We first calculated the distance between individual cells according to the cosine similarity and then constructed a KNN network with *K* = 40 (for details see Methods). We applied the *Louvain* algorithm^52^, detected communities of cells, and used the *F - measure* (the harmonic mean between precision and sensitivity) to check the degree to which the inferred communities reproduce the known cell types from the original publication. We found that NBC resolves the original seven cell types with high precision and sensitivity (Fig 2a and b, *F - measure* = 0.93). Constructing the network with *K* = 10 resulted in a slightly lower *F - measure* (Fig 2c and d, *F - measure* = 0.81), mainly due to the separation of one original cluster (indicated by light blue in C) into two inferred clusters (indicated by light blue and orange in D). Evaluating the precision and sensitivity of NBC for a wide range of K’s shows that, for this dataset, NBC is quite robust to the choice of *K* for values larger than *K* ≈ 18 (Fig 2e). Using the correlation similarity for constructing the KNN network results in a similar performance in terms of the *F - measure* (Fig 2e). In this dataset, NBC outperformed other common clustering methods (Table 2).

**Figure 2.**
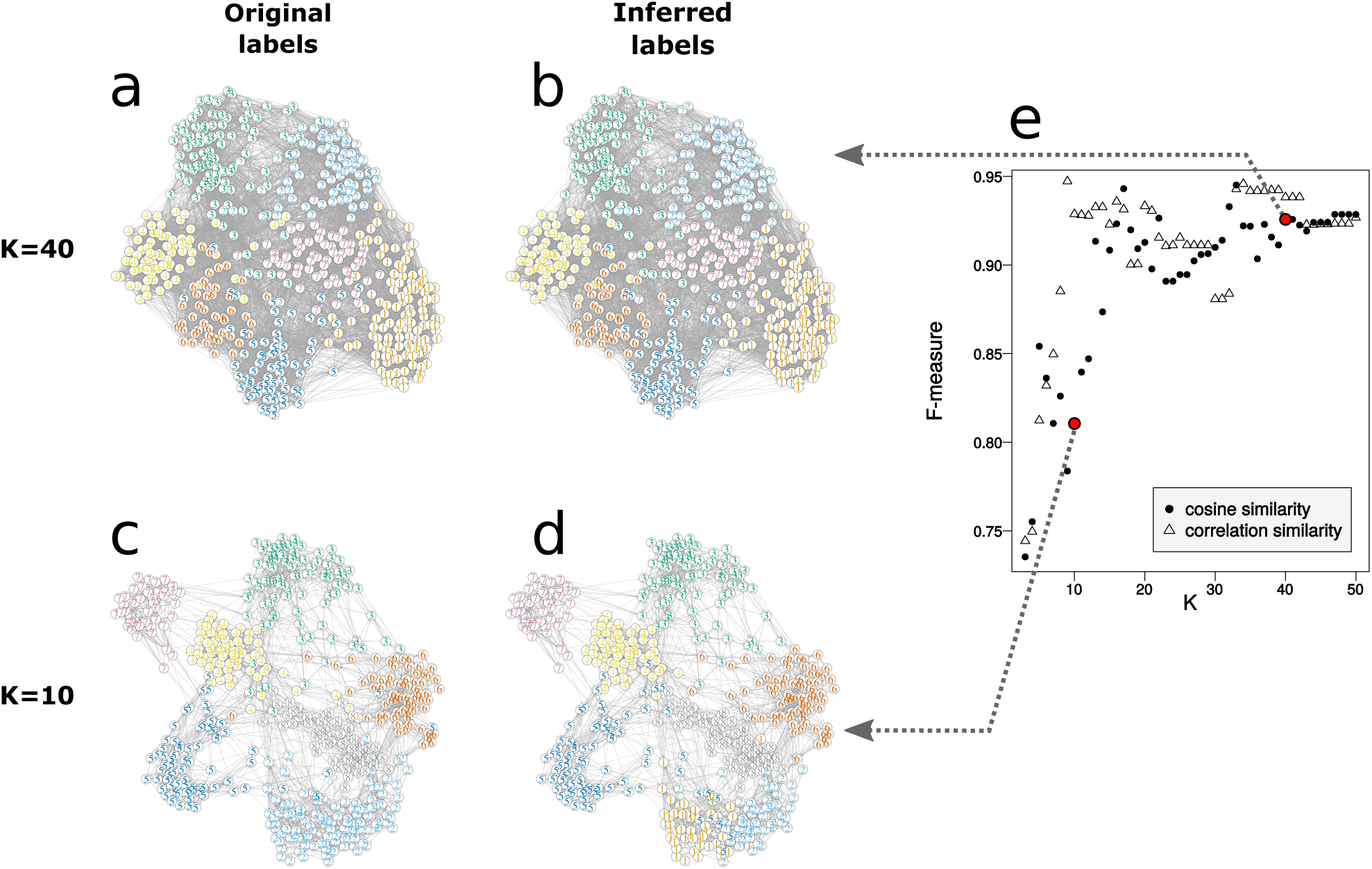
NBC accurately resolves seven cell types from a glioblastoma single-cell RNA-seq dataset. We applied NBC to single-cell RNA-seq data published by Patel *et al.*^62^ *containing single cells collected from five patients with glioblastoma and two gliomasphere cell lines. A KNN network constructed using cosine similarity with K* = 40 is shown in (a) and (b). A similar KNN network with *K* = 10 is shown in (c) and (d). Nodes shown in (a) and (c) are color-coded according to cell types reported by the original publication, while nodes in (b) and (d) are color-coded according to communities inferred by the *Louvain* algorithm. (e) The *F-measure*, which is the harmonic mean of precision and recall, is plotted against *K* for networks constructed with cosine (full circles) and correlation (empty triangles) similarities. Specific points corresponding to *K* = 10 and *K* = 40 are highlighted in red.

**Table 2.** Comparison between NBC, K-means, hierarchical clustering, and spectral clustering in terms of the *F-measure* for two single-cell RNAseq datasets. The *F-measure*, which is the harmonic mean between precision and sensitivity, measures the degree to which the inferred communities reproduce the known cell types from the original publication.

| Method | Patel et al.<sup>62</sup> dataset<br>(Fig 2) | Klein et al.<sup>73</sup> dataset<br>(Fig 3) |
| --- | --- | --- |
| NBC<br>(Cosine similarity, *Louvain* CD) | 0.93 ($K = 40$) | 0.81 ($K = 30$) |
| NBC<br>(Correlation similarity, *Louvain* CD) | 0.94 ($K = 40$) | 0.90 ($K = 30$) |
| K-means (Euclidean similarity) | 0.76 | 0.84 |
| Hierarchical clustering<br>(Euclidean similarity) | 0.86 | 0.44 |
| Hierarchical clustering<br>(Correlation similarity) | 0.79 | 0.44 |
| Spectral clustering | 0.47 | 0.80 |

We performed a similar analysis on another single-cell RNA-seq dataset published by Klein *et al.*^73^ containing 2,717 mouse embryonic stem cells collected from four consecutive developmental stages. We found that also for this dataset, NBC performs well relative to other common clustering methods (Fig 3 and Table 2). However, here we found that, for sufficiently large *K* (*K >* 50), correlation similarity had better performance than cosine similarity (*F - measure* = 0.90 for correlation similarity and 0.81 for cosine similarity, in both cases using the *Louvain* algorithm, Fig 3e and Table 2).

**Figure 3.**
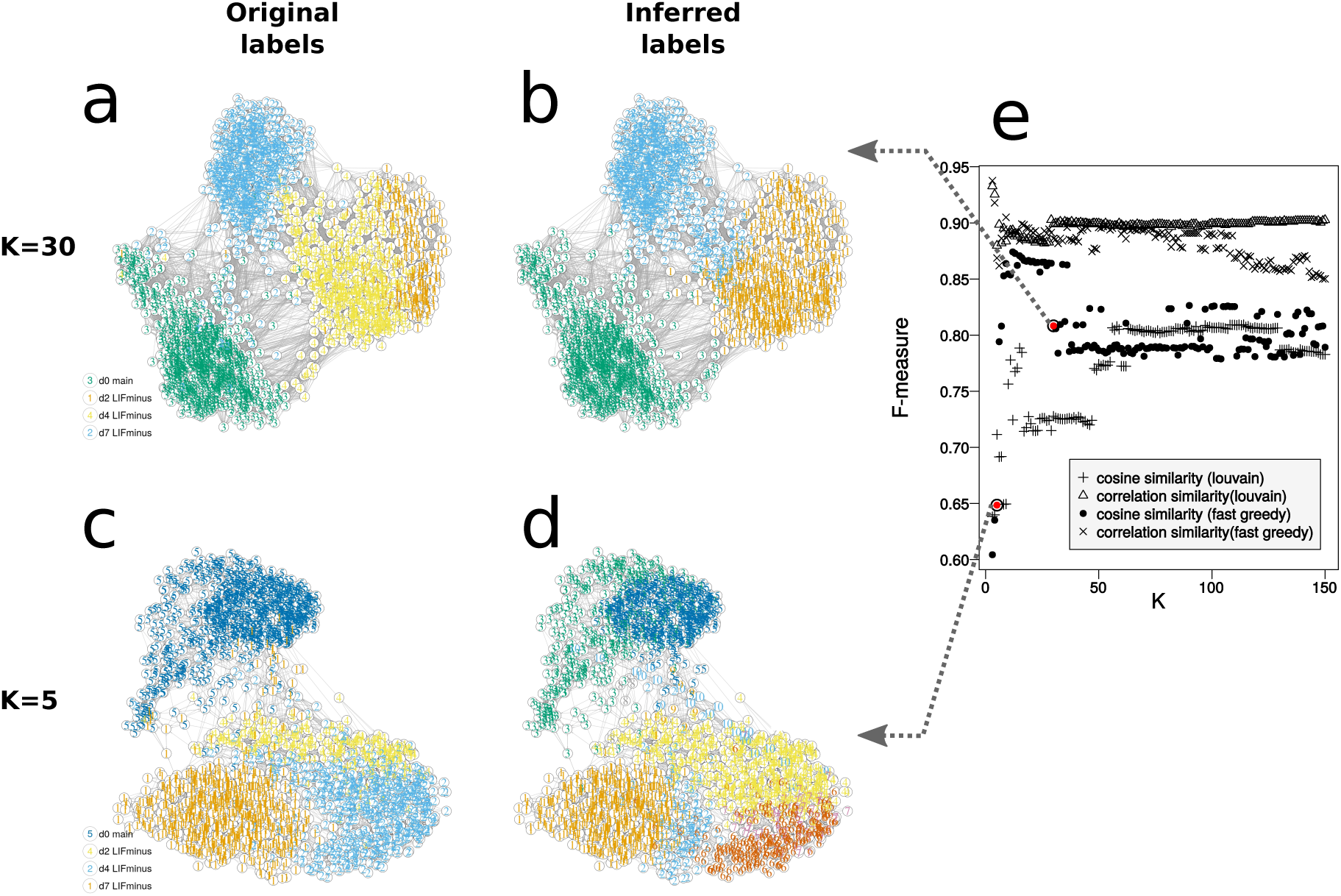
NBC resolves four differentiation stages in a single-cell RNA-seq dataset from mouse embryonic stem cells. We applied NBC to single-cell RNA-seq data published by Klein *et al.*^73^ *containing single mouse embryonic stem cells collected from four consecutive developmental stages. A KNN network constructed using cosine similarity with K* = 30 is shown in (a) and (b). A similar KNN network with *K* = 5 is shown in (c) and (d). Nodes shown in (a) and (c) are color-coded according to the differentiation stage reported by the original publication, while nodes in (b) and (d) are color-coded according to communities inferred by the *fast greedy* algorithm. (e) The *F-measure*, which is the harmonic mean of precision and recall, is plotted against *K* for networks constructed with cosine and correlation similarities and communities inferred by the *fast greedy* and *Louvain* algorithms. Specific points corresponding to *K* = 5 and *K* = 30 are highlighted in red.

### Comparing NBC with other common clustering methods

We compared NBC with three widely used clustering algorithms: K-means, hierarchical clustering, and spectral clustering. As a reference dataset, we used 3,174 human tissue-specific gene expression samples from the GTEx project^1^ that were collected from 21 tissue types. Based on the number of tissue types in the original samples, the number of clusters was set to 21 for all clustering algorithms (see Methods). A KNN network was constructed with *K* = 50 and the *Louvain* algorithm was applied to infer sample communities. In order to compare the algorithms and test their robustness to different levels of “noise”, a fraction of the genes was randomly permuted by shuffling their sample labels (Fig 4a), thereby maintaining the original distribution for each gene. This actually mimics different levels of non-informative features. We observed that for this data set, NBC out-performs K-means, spectral, and hierarchical clustering in terms of the *F-measure* over a wide range of noise levels (Fig 4a).

**Figure 4.**
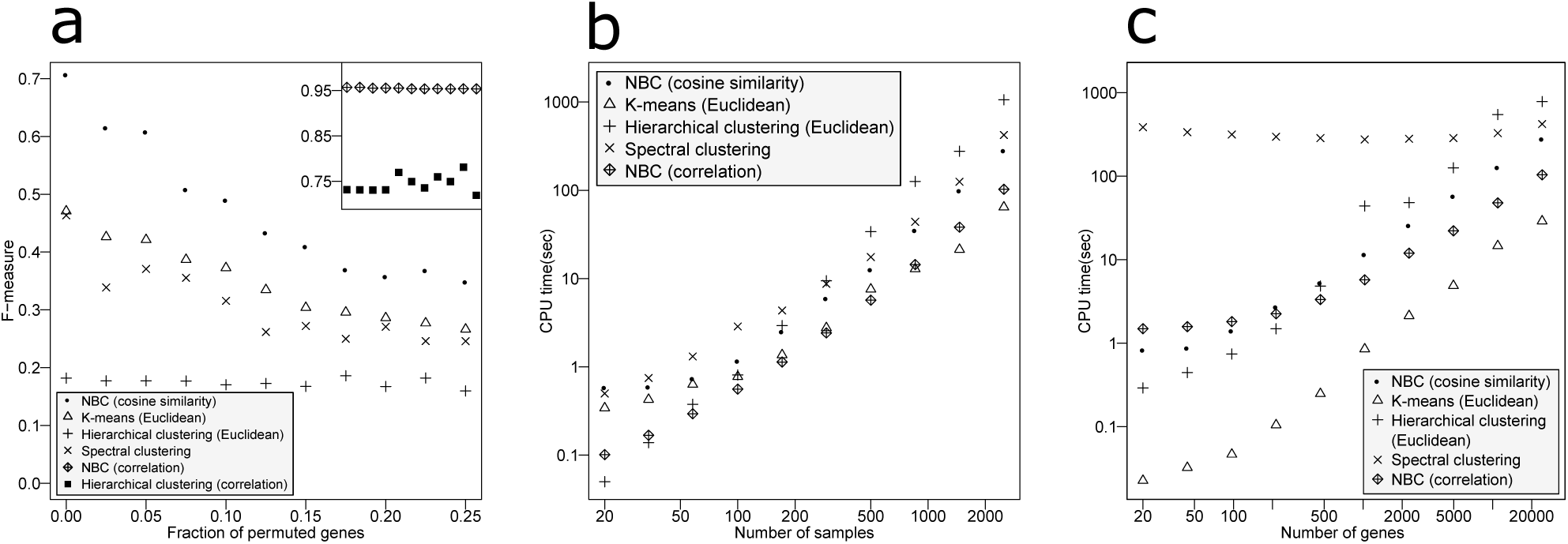
Comparing NBC with other common clustering methods. (a) Shown is a comparison between NBC, K-means, hierarchical clustering, and spectral clustering in terms of the *F-measure* as a function of the effective number of genes. We effectively reduced the number of genes by randomly permuting the sample labels for a fraction of the genes. The inset shows results for NBC and hierarchical clustering with correlation similarity. (b,c) Shown is a comparison of the CPU times required by the five clustering methods, once as a function of the number of randomly chosen samples while keeping the number of genes fixed to 24,071 (b), and once as a function of the number of randomly chosen genes while keeping the number of samples fixed to 2,500 (c). Each point shown is an average of three iterations. Data used in (a) was downloaded from the GTEx consortium^1^, and data used in (b) and (c) was taken from a single-cell RNA-seq dataset published by Macosko et al.^13^.

We performed a similar comparison of the CPU time required to detect communities using a single-cell RNA-seq dataset that was published by Macosko et al.^13^. We randomly chose subsets of varying sizes of samples (cells) and genes in order to test the dependency of the running-time on the size of the dataset. We found that NBC falls in-between K-means, Hierarchical clustering, and spectral clustering for a wide range of samples and genes (Fig 4b and c).

### NBC can be used to resolve tissue-specific genes

NBC can also be used to detect communities of genes from large gene expression datasets. To demonstrate this, we analyzed a dataset composed of 394 GTEx samples collected from three tissues: pancreas, liver, and spleen. First, a KNN network with *K* = 10 was constructed for the 394 samples. The resulting network contained three perfectly separated components, each corresponding to one of the three tissue types (Fig 5b-d, *F - measure* = 1). We then constructed another KNN network for the 27,838 genes with *K* = 200. The *Louvain* community detection algorithm was applied and 11 gene communities were detected.

**Figure 5.**
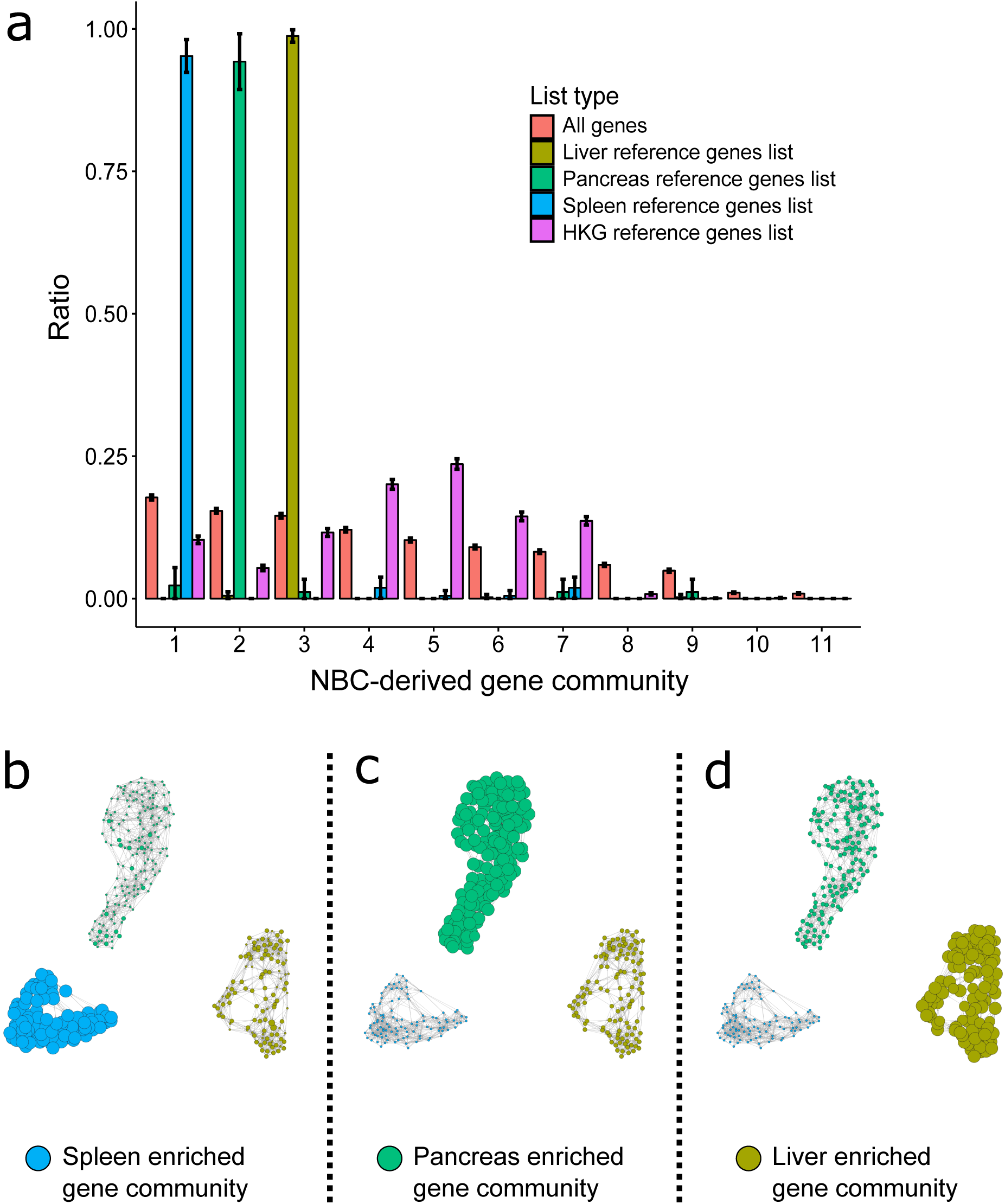
NBC can be used to resolve tissue-specific genes. (a) NBC was applied to a mix of RNA-seq expression profiles from “bulk” samples of the human liver, pancreas, and spleen, that were obtained from the GTEx project^1^. Eleven communities of genes were detected by NBC using the *Louvain* algorithm. For each NBC-derived gene community, the relative fractions of all genes, housekeeping genes (HKG), and three tissue-specific reference genes lists are shown. The tissue-specific reference genes lists were downloaded from the Human Protein Atlas^74^. The NBC-derived gene communities are ordered according to their relative sizes, which is represented by the fraction of total genes that belong to each community (light red bars). (b-d) Shown is the network of samples, where in each panel the node size is proportional to the average log-transformed expression of the genes from NBC-derived community #1 (spleen-enriched, panel b), NBC-derived community #2 (pancreas-enriched, panel c), and NBC-derived community #3 (liver-enriched, panel d). The nodes are color-coded according to their respective tissue type.

In order to explore the biological meaning of these 11 NBC-derived gene communities, we used three independently derived tissue-specific “reference lists” of genes from the Human Protein Atlas^74^ that were found to be over-expressed in the pancreas, liver, and spleen. We compared these three reference lists to the 11 NBC-derived communities and found that each reference list was found predominantly in a single community (Fig 5a). In community #1, 200 of the 210 spleen-specific reference genes were found, in community #2, 82 of the 87 pancreas-specific reference genes were found, and in community #3, 397 of the 403 liver-specific reference genes were found. On the contrary, a reference list of “House Keeping Genes” (HKG) was found to be distributed relatively uniformly among the different communities.

Another helpful feature of NBC is that it can be used to visualize families of genes within the network of samples. To demonstrate this, we measured the relative gene expression levels of genes from NBC-derived community no. 1 (enriched for spleen related genes) in each node (=sample) of the network and used this information to determine the size of that node (Fig 5b). It can be seen that the average expression level of NBC-derived community #1, that is enriched for spleen specific genes, was indeed much higher in the spleen samples compared to pancreas and liver samples (Table 3, one side t-test p-value=2 *×* 10^−61^). We observed similar results when we repeated this analysis for NBC-derived communities 2 and 3 (pancreas-enriched and liver-enriched, Fig 5c and d, Table 3).

**Table 3.** Summary statistics for NBC-derived gene communities.

| NBC-derived gene community no. | Total number of genes in community | Annotation of enriched tissue-specific "reference" genes list (from Human Protein Atlas) | Enrichment - number of genes from tissue-specific "reference" list (from the Human Protein Atlas) found in this NBC-derived community | p-value for over-expression in corresponding tissue-specific samples (one side t-test with unequal variance) |
| --- | --- | --- | --- | --- |
| 1 | 4940 | Spleen | 200 of 210 | $2 \times 10^{-61}$ |
| 2 | 4284 | Pancreas | 82 of 87 | $2 \times 10^{-250}$ |
| 3 | 4043 | Liver | 397 of 403 | $2 \times 10^{-124}$ |

## Discussion

To date, genomic datasets typically contain hundreds of samples with thousands of features each. FACS datasets may contain millions of samples with 10-30 features each. Improvements in single-cell processing techniques (e.g. droplet-based^13^ or magnetic-beads based methods^75^) are further increasing the number of samples in single-cell RNA-seq data. Therefore, tools for genomic data analysis need to perform efficiently on very large datasets. In this aspect, the major bottleneck of our toolkit is the KNN network construction step for which we use the ball tree algorithm^63^. Although efficient, this algorithm does not scale well with respect to memory usage and query time when the number of features increases. One possible solution is to use methods for approximating KNN networks, which might be utilized for this step after careful exploration of the error they introduce^76, 77^. Another possibility is to use parallel architectures to accelerate KNN network construction^78^.

Moreover, when the number of features *P* is larger than the number of samples *N* (*P > N*), network construction can be made more efficiently by projecting the original matrix (embedded in ℝ^*P*^) into a lower dimension space (ℝ^*N*^) using PCA^79^. This projection preserves the original Euclidean distances between the samples. Other possible projections into lower dimensions are truncated SVD or approximated PCA methods^80^. However, these do not preserve the original Euclidean distances between the samples^80^. The original matrix can also be projected into lower dimensions by non-linear transformations like tSNE^81^. tSNE captures much of the local structure of the data and has been widely used for single-cell expression analysis^21, 82^.

Note that Network-based methods themselves can be used for dimensionality reduction. Isomap^83^, for instance, constructs a KNN graph that is used to approximate the geodesic distance between data points. Then, a multi-dimensional scaling is applied, based on the graph distances, to produce a low-dimensional mapping of the data that maintains the geodesic distances between all points.

NBC has much in common with the widely used density-based clustering method DBSCAN^33^. Although both methods explore the local structure of the data, NBC uses the *K* nearest neighbors, while DBSCAN defines clusters by their local densities. However in NBC, as opposed to DBSCAN, no minimal distance is required to define two samples as neighbors. In addition, DBSCAN does not produce the network explicitly, but rather just the connectivity component of each sample. This is in contrast to NBC that provides an explicit representation of the underlying weighted network that can be analyzed with different CD algorithms.

NBC requires the user to specify the following parameters: a similarity measure, a community detection algorithm, and the number of nearest neighbors *K*. For the datasets that we checked we found that NBC is not very sensitive to the choice of *K* given sufficiently large values (*K >* 18) (e.g. Fig 2e); however, we found that the choice of the community detection algorithm and especially the similarity measure may significantly influence its performance (e.g. Fig 3e and Fig 4a). Hence, these parameters should be chosen carefully when applying NBC to other data types. Similar to other machine learning approaches, NBC parameters can be optimized using a labeled training dataset prior to application on unlabeled data.

We created an open and flexible python-based toolkit for Networks Based Clustering (NBC) that enables easy and accessible KNN network construction followed by community detection for clustering large biological datasets, and used this toolkit to test the performance of NBC on previously published single-cell and bulk RNA-seq datasets. We find that NBC can identify communities of samples (e.g. cells) and genes, and that it performs better than other common clustering algorithms over a wide range of parameters.

In practice, given a new dataset, we recommend to carefully test different alternatives for network construction and community detection since results may vary among different datasets according to their unique characteristics. We believe that the open and flexible toolkit that we introduced here can assist in rapid testing of the many possibilities.

## Methods

### single-cell and “bulk” RNA sequencing datasets

We used Four datasets in this study:

I. Single-cell RNA-seq data from Patel et al.^62^ containing single-cell gene expression levels from five patients with glioblastoma and two gliomasphere cell lines that were acquired using the SMART-SEQ protocol. We downloaded the preprocessed data from GEO^84^. Altogether, this dataset contains 543 cells by 5,948 genes.
II. Single-cell RNA-seq data from Klein et al.^73^ containing single-cell gene expression levels from mouse embryonic stem cells at different stages of differentiation that were acquired using the inDrop protocol. In that experiment, samples were collected along the differentiation timeline by sequencing single cells at 0, 2, 4, 7 days after withdrawal of leukemia inhibitory factor (LIF). We downloaded the preprocessed data from GEO^85^ and removed genes with zero expression levels, resulting in a dataset of 8,669 cells by 24,049 genes. For the analysis presented in Fig 3, we first removed technical replicates and data from a control cell line, resulting in a total number of 2,717 cells.
III. “Bulk” RNA sequencing datasets from the Genotype-Tissue Expression (GTEx) database^1^. We downloaded the data from the GTEx website^86^ version 6. This dataset includes 8,555 samples taken from 30 tissue types (according to the SMTS variable) of 570 individuals. Gene names were translated from Ensemble gene ID into HGNC symbols using the *BioMart* Bioconductor package^87^. In cases where we found multiple matches of the Ensemble gene ID’s corresponding to a single HGNC symbol, the Ensemble gene ID with maximum average intensity across all samples was chosen. To compare different clustering methods (Fig 4a) we chose only samples originating from a single tissue type. Moreover, we omitted tissues having multiple detailed tissue types (according to the SMTSD variable) even if they had a single tissue type (indicated by the SMTS variable). Likewise, genes with zero expression were omitted, resulting in a dataset of 3,174 samples by 33,183 genes from 21 tissue types.
IV. Single-cell RNA-seq data from Macosko et al.^13^ containing single-cell gene expression levels from a P14 mouse retina that were acquired using the Drop-Seq protocol. We downloaded the preprocessed data from GEO^88^ and removed genes with zero expression, resulting in a dataset of 49,300 cells by 24,071 genes. This dataset was used to compare the performance, in terms of CPU time, of NBC, K-means, and hierarchical clustering as shown in Fig 4b-c.

### KNN network construction and visualization

A KNN network with cosine similarity was constructed using the *scikit-learn* package^89^ for machine learning in Python. Since the cosine distance was not directly available in the *scikit-learn* BallTree() function, we used a two-step implementation as follows: First, each sample was mean-centered and standardized such that it will have zero mean and unit length (L2 normalization). Next, the ball tree algorithm^63^ was applied with Euclidean distance to find the *K* nearest neighbors of each sample and construct a KNN network. Then, the Euclidean distances between the nodes (=samples) were transformed to cosine similarities that were used as the edges weights for community detection.

We calculated the cosine similarity from the Euclidean distance as follows. The cosine similarity between two vectors A and B is defined as:

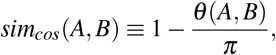

where *θ* (*A, B*) is the angle between A and B. If A and B are also of unit length (L2 normalized) then this angle is related to the Euclidean distance *D*_*euc*_(*A, B*) according to:

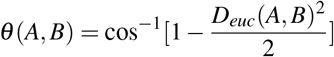

or: 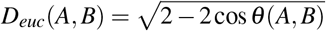.

Network layouts for visualization were created by the *fruchterman-reingold* algorithm^90^ as implemented in the *igraph* Python and R packages^91^. For correlation similarity we calculated the full *spearman* correlation matrix *ρ*(*A, B*) between any two vectors *A* and *B* using the *corr* function in R.

### Community detection algorithms

In this manuscript we generally used the *Louvain*^52^ *algorithm for community detection as implemented by the igraph* Python and R packages for network analysis^91^, apart from Fig 3 in which we used the *fast greedy*^47^ *algorithm.*

*The Louvain* method partitions the nodes of the network into communities *c*1, *c*2, *c*3,*…*, such that network modularity score

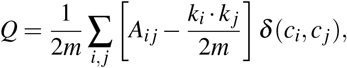

is maximized. In the above formula, *A*_*ij*_ is the edge weight between nodes *i* and *j, k*_*i*_ is the degree of node *i* (that is, the sum of the weights of all the links emanating from node *i*), *m* is the overall sum of weights,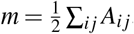, and *δ* (*c*_*i*_, *c* _*j*_) is the Kronecker delta function. The network modularity score is actually the difference between the number of links connecting nodes within the same community and the expected number of links in a network with randomly shuffled links.

Briefly, the algorithm starts by assigning a separate community to each node. Then, the algorithm iterates between two steps: In the first step, the modularity is maximized by repeatedly iterating over all nodes in the network. For each node, we evaluate the gain in modularity that will take place by removing it from its present community and assigning it to one of its neighboring communities. If the overall modularity can be improved, the community of the node is reassigned accordingly. This process is repeated until a local maximum is reached. In the second step, the algorithm constructs a meta-network in which the nodes are communities from the first step and the edges are the edges between the communities. At this point, the first step is repeated on the nodes of the new meta-network in order to check if they can be merged into even larger communities. The algorithm stops when there is no more improvement in the modularity score.

### Statistical measures for comparing NBC and other common clustering algorithms

To compare the performance of NBC, hierarchical clustering, K-means, and spectral clustering, we used the *F-measure*, which is the harmonic mean between the precision *P* and sensitivity (recall) *R*:

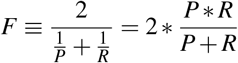

where 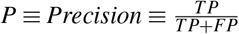, and 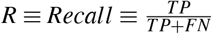 (*TP*-true positive, *FP*-false positive, *FN*-false negative). To calculate precision and sensitivity for each clustering algorithm, we also used the R package *clusterCrit*^92^ that compares the labels from the original publication to the labels inferred by the algorithm.

Another requirement for evaluating and comparing the different clustering algorithms was to require all of them to find the same number of clusters. Therefore, the number of required clusters was set to the number of distinct groups from the original publication (7 clusters in Fig 2, 4 clusters in Fig 3, 21 clusters in Fig 4a, etc.). We used the stat package in R^93^ to run hierarchical clustering and K-means clustering with default parameters (Euclidean distance), apart from the linkage in hierarchical clustering which was set to *average* linkage. For spectral clustering we used the *specc* function from the *kernlab* R package^94^ with default parameters. Generally, all parameters were chosen as default unless otherwise specified.

All computations were done on a standard PC with i7-4600 CPU with 2.10 GHz and 16 GB of RAM memory.

### Tissue-specific reference genes lists from the Human Protein Atlas

Tissue-specific reference lists of genes were obtained from the Human Protein Atlas^74^ version 14^95^. Altogether, the house-keeping genes (HKG) reference list is composed of 8,588 genes, and the liver-specific, pancreas-specific, and spleen-specific reference genes lists are composed of 436, 234, and 95 genes respectively. Genes that do not appear in the dataset or genes that appear in more than one tissue-specific list were removed, resulting 8331, 403, 210, and 87 genes in the HKG, liver-specific, pancreas-specific, and spleen-specific reference genes lists respectively.

