## Supplementary material for "Applications of community detection algorithms to large biological datasets": Ipython notebook (from another version of this preprint: version 1, 2019-02-12)

Supplementary\_IPython\_notebook


We created an open and flexible python-based toolkit for Networks Based Clustering (NBC) that enables easy and accessible network construction and community detection.

This toolkit allows the user to follow and modify some aspects of the algorithm, that is, to map a given dataset into a KNN network, to visualize the network, and to perform community detection using a variety of similarity measures and community detection algorithms of his choice.

### Prerequisites and required packages:¶

Before starting, install and load the following packages.
  
  
Notes about the python-igraph package:

1. The python-igraph package requires the python cairo (pycairo) package.
2. iGraph has a known bug that prevents iPython/Jupyter from plotting graphs using the plot() function (see
   https://github.com/igraph/python-igraph/issues/88).
   This can be fixed manually by downloading and replacing the local file ".../igraph/drawing/\_\_init\_\_.py" with a fixed file from github:
   https://raw.githubusercontent.com/igraph/python-igraph/8864b46849b031a3013764d03e167222963c0f5d/igraph/drawing/\_\_init\_\_.py

In [1]:

```
#%% Import python modules:
import numpy as np
from sklearn.neighbors import BallTree
from sklearn.metrics import f1_score
from sklearn.preprocessing import scale
from sklearn.decomposition import PCA
from sklearn.manifold import TSNE
from collections import Counter
from igraph import *
import matplotlib.pyplot as plt
import seaborn as sns
import itertools
from sklearn import preprocessing
import urllib.request as ur
from gzip import GzipFile
import pandas as pd
```

In [2]:

```
#%% Initialize:
np.random.seed(1)
%matplotlib inline
```

### Data retrieval and exploration:¶

First, the single cell RNA-seq dataset from ***"Single-cell RNA-seq highlights intratumoral heterogeneity in primary glioblastoma"*** by Patel et al. was retrieved and pre-processed (re-naming, removing bulk samples, and scaling).

In [3]:

```
#%% Download the Patel et al. dataset from GEO:
url = ('ftp://ftp.ncbi.nlm.nih.gov/geo/series/' 'GSE57nnn/GSE57872/suppl/GSE57872_GBM_data_matrix.txt.gz')
req = ur.Request(url) 
z_f = ur.urlopen(req)
f = GzipFile(fileobj=z_f, mode="r")
df = pd.read_csv(f, sep='\t')
df.drop(df.columns[0], axis=1,inplace=True)
import re
cellLabels = [re.sub('(.*)_.*','\\1',x) for x in list(df.columns.values)]
# Correct a typo in the cell names:
cellLabels = [w.replace('MGH264','MGH26') for w in cellLabels]
# Removing bulk samples (omit groups that are actually composed of one sample):
from collections import Counter
multiLabels = [k for k, v in Counter(cellLabels).items() if v > 1 ]
matching = [i for i, x in enumerate(cellLabels) if  x in multiLabels] # Removing bulk samples
dataMat = scale(df.iloc[:,matching].values, axis=0)               # Scaling
cellLabels = [cellLabels[i] for i in matching]
```

The dataset contains expression levels of 5948 genes (rows) over 532 single cells (columns):

In [4]:

```
#%% The gene expression matrix:
dataMat.shape
```

Out[4]:

```
(5948, 532)
```

The dataset contains 7 cell sets: glioblastoma tumors from five individuals (MGH) and two gliomasphere cells lines (CSC):

In [5]:

```
#%% The cell labels:
Counter(cellLabels)
```

Out[5]:

```
Counter({'MGH26': 118,
         'MGH28': 94,
         'MGH29': 75,
         'MGH30': 73,
         'MGH31': 70,
         'CSC6': 44,
         'CSC8': 58})
```

Exploratory data analysis was performed using PCA and tSNE which are methods for dimension reduction.

#### PCA:¶

The first two principal components are not sufficient for separating the samples.

In [6]:

```
#%% PCA:
pca = PCA(n_components=2)
X_r = pca.fit_transform(dataMat.T )

plt.figure()
colors = sns.color_palette("hls", list(set(cellLabels)).__len__())
lw = 2
for color,  target_name in zip(colors, list(set(cellLabels))):
    IDX = [i for i,x in enumerate(cellLabels) if x==target_name]
    plt.scatter(X_r[IDX, 0], X_r[IDX, 1], color=color, alpha=.8, lw=lw,
                label=target_name)
plt.legend(loc=2, scatterpoints=1,bbox_to_anchor=(1.05, 1))
plt.xlabel('PC1')
plt.ylabel('PC2')
plt.title('PCA');
```

#### tSNE:¶

In contrast, the two first components of tSNE were better able to separate the samples.

In [7]:

```
#%% tSNE:
tsne = TSNE(n_components=2, init='pca', random_state=0)
X_r = tsne.fit_transform(dataMat.T)

plt.figure()
colors = sns.color_palette("hls", list(set(cellLabels)).__len__())
lw = 2
IDX = [i for i,x in enumerate(cellLabels) if x==target_name]
for color,  target_name in zip(colors, list(set(cellLabels))):
    IDX = [i for i,x in enumerate(cellLabels) if x==target_name]
    plt.scatter(X_r[IDX, 0], X_r[IDX, 1], color=color, alpha=.8, lw=lw,
                label=target_name)
plt.legend(loc=2, scatterpoints=1,bbox_to_anchor=(1.05, 1))
plt.xlabel('DIM1')
plt.ylabel('DIM2')
plt.title('t-SNE');
```

### Construction of a weighted KNN network:¶

A KNN network with cosine similarity was constructed using K=50 nearest neighbors.

Since the cosine distance was not directly available in the *scikit-learn* BallTree() function, we used a two-step implementation as follows:

First, each sample was mean-centered and standardized such that it will have zero mean and unit length (L2 normalization).

Next, the ball tree algorithm was applied with Euclidean distance to find the $K$ nearest neighbors of each sample and construct a KNN network.

Then, the Euclidean distances between the nodes (=samples) were transformed to cosine similarities that were used as the edges weights for community detection.

In [8]:

```
#%% First, calculate distances and find the K nearest neighbors for each sample:
# The scikit-learn package expects a [n_samples, n_features] array.
dataMat_T=dataMat.T  

# L2 normalization: scale each sample vector such that it will have zero mean and unit length:
dataMat_T = scale(dataMat_T,axis=1)/np.sqrt(dataMat_T.shape[1]) 

# For each sample, find the K nearest neighbors according to their Euclidean distances:
bt = BallTree(dataMat_T, metric = 'euclidean')
#
# Additional metrics are supported:
#bt = BallTree(dataMat_T, metric = 'manhattan') # Manhattan distance metric
#bt = BallTree(dataMat_T, metric = 'chebyshev') # Chebyshev distance metric

dist, ind = bt.query(dataMat_T, k=50, return_distance=True)
```

We calculated the cosine similarity from the Euclidean distance as follows.
  
The cosine similarity between two vectors A and B is defined as:
$$
sim\_{cos}(A,B)\equiv1-\frac{\theta(A,B)}{\pi},
$$
where $\theta(A,B)$ is the angle between A and B.
  
If A and B are also of unit length (L2 normalized) then this angle is related to the Euclidean distance $D\_{euc}(A,B)$ according to:
$$
\theta(A,B)=\cos^{-1}[1-\frac{D\_{euc}(A,B)^2}{2}]
$$
or: ${D\_{euc}(A,B)=\sqrt{2-2\cos\theta(A,B)}}$.

In [9]:

```
#%% Calculate similarities and construct a weighted graph:
# Initalize:
vertices = np.arange(dataMat_T.shape[0])
edges = [(i,ind[i,j]) for i in range(ind.shape[0])
         for j in range(1,ind.shape[1]) ]
dist = dist[:,1:].flatten()

# Transform Euclidean distances to cosine similarities:
weights = 1 - np.arccos(1-(dist**2)/2)/np.pi
#
#weights = 1/(1+dist)      # Change distance to similarity measure
#weights = 1/np.exp(dist)

# Assign Weights to edges and create a weighted graph:
g = Graph(edges, edge_attrs=dict(weight=weights,directed=True))
```

We assign colors to the nodes of the graph according to the original cell labels:

In [10]:

```
#%% Assign colors to the nodes of the graph according to the original cell labels:
_, indices = np.unique(np.array(cellLabels), return_inverse=True)
col_palette_i = sns.color_palette("husl", max(indices)+1)
g.vs["color"] = [col_palette_i[i] for i in indices]
```

### Network visualization:¶

The network was visualized using the force-directed Fruchterman Reingold layout. Nodes were color-coded according to their cell types.

In [11]:

```
#%% Network visualization:
layout = g.layout("fruchterman_reingold")
#
#Additional layouts are supported:
#layout = g.layout("kamada_kawai") # Kamada_kawai layout
#layout = g.layout("lgl")          # Large Graph Layout
#....

plot(g, layout = layout, edge_width = 0.15, vertex_size = 9)
```

Out[11]:

xml version="1.0" encoding="UTF-8"?

### Community detection:¶

We used the *Louvain* algorithm for community detection. Nodes were color-coded according to the inferred communities.

In [12]:

```
#%% Community detection:
communities = g.community_multilevel(weights = g.es["weight"])  # The Louvain algorithm (Blondel et al.)
#
# Additional community detection (CD) algorithms are supported: (http://igraph.org/python/doc/igraph.Graph-class.html):
#communities = g.community_fastgreedy(weights = g.es["weight"]) # fastgreedy CD 
#communities = g.community_spinglass(weights = g.es["weight"])  # spinglass CD (Reichardt et al.)
#communities = g.community_infomap(weights = g.es["weight"])    # infomap CD (Rosvall et al.)
#...

# Reassign colors to the nodes of the graph according to the inferred communities:
ms = communities.membership
col_palette = sns.color_palette("husl", max(ms)+1)
g.vs["color"] = [col_palette[i] for i in ms]

plot(g, layout = layout, edge_width = 0.15, vertex_size = 9)
```

Out[12]:

xml version="1.0" encoding="UTF-8"?

### Statistical assessment of community detection:¶

We used the *F-measure*, which is the harmonic mean between the precision $P$ and sensitivity (recall) $R$, to check the degree to which the inferred communities reproduce the known cell types from the original publication:
$$
F= \frac {2} {\frac{1}{P}+\frac{1}{R}}=\frac {2P\*R}{P+R}
$$
Where:
$$
P \equiv Precision=\frac {TP}{TP+FP}
$$
  
$$
R \equiv Recall=\frac {TP}{TP+FN}
$$
  
(Note that it is possible to calculate the precision and sensitivity also with the ***clusterCrit*** R package that compares the cell labels from the original publication to the labels inferred by the community detection algorithm).

In [13]:

```
#%% Statistical assessment community detection:
org_paired = indices[:,None]==indices
com_paired = np.asarray(ms)[:,None]==np.asarray(ms)
TP = np.logical_and(org_paired,com_paired).sum()
FP = np.logical_and(np.logical_not(org_paired),com_paired).sum()
FN = np.logical_and(org_paired,np.logical_not(com_paired)).sum()
F_measure = 2*TP/(2*TP+FP+FN)
F_measure
```

Out[13]:

```
0.9294139160647343
```
